# Evaluation of diversity levels of the integrase gene sequences coming from HIV-1 virus, supporting the lack of target specificity of ivermectin *versus* the integrase-importin complex in SARS-CoV-2 infection

**DOI:** 10.1101/2020.07.18.210096

**Authors:** Pierre Teodósio Félix

## Abstract

Therapies with new drugs have been appearing in tests worldwide as potential inhibitors of sars-cov-2 virus replication. Recently, one of these drugs, Ivermectin, was reported as an inhibitor of the nuclear import of HIV-1 proteins *in vitro*, soon becoming the target of an international prospecting work (not yet published), with patients tested for COVID-19. However, understanding the evolutionary aspects of the biological components involved in the complex drug-nuclear import helps in understanding how these relationships exist in the deactivation of viral infections. Thus, 153 sequences of the HIV-1 integrase gene were analyzed for their genetic structure and molecular diversity and the presence of two distinct groups for the Gene and not only one, was detected; As well as different degrees of structuring for each of these groups. These results support the interpretation of the lack of conservation of the HIV-1 gene and that the number of existing polymorphisms, only for this structure of the complex, implies the non-efficiency of a drug at population levels. Thus, the molecular diversity found in HIV-1 can be extrapolated to other viruses, such as Including, SARS-CoV-2 and the functionality of the drug, interacting with the integrase-importin complex, can be further decreased.

## 1. Introduction

As a flattening measure of the growth curve of the number of cases of COVID-19 in Brazil, the recommendations of the Ministry of Health continue to be limited to the monitoring and containment of the virus as well as to those of maintaining social distancing, the use of face protection masks and constant hand washing. These recommendations, which follow the recommendations of all the world’s health agencies, seem to be the most effective to inhibit the outbreak of this pandemic, which, as a direct consequence of non-control, would result in the breakdown of our health system (MINISTÉRIO DA SAÚDE, 2020).

However, a number of therapies are in tests worldwide and these range from vaccines to the use of some drugs (WHO, 2020). With regard to the new drugs tested to combat COVID-19, a not-so-new one (for other treatments) called IVERMECTIN, figured as an alternative for its power to inhibit the replication of the SARS-CoV-2 virus *in vitro*. Ivermectin, a pre-tested food and drug administration (FDA) for antiparasitic use, demonstrated broad-spectrum antiviral activity *in vitro* for the SARS-CoV-2 virus, and even two hours after infection was able to reduce the amount of viral RNA by approximately 5,000 times after 48 hours. This made ivermectin a candidate for in-depth investigations for possible human benefits (CALY, L. *et al*., 2020).

Even thinking about the “*in vitro*” characteristics of the study with ivermectin, its usefulness and potentiality as therapy did not reach exhaustion. Contrary to some drugs such as Chloroquine and hydroxichloroquine, discarded by who and many health agencies and research centers around the world, it ended up becoming a target in an international, multicenter and observational prospecting work, controlled on a case-by-case basis, using data collected from patients diagnosed with COVID-19 between January 1 and March 31, 2020. These patients were exposed to doses of Ivermectin compared to patients with COVID-19 who received medical treatment without ivermectin. In this study (*in vivo*) and not yet published, the researchers assume that, in addition to being safe for use, the administration of ivermectin in patients hospitalized with COVID-19 was directly associated with the fact that a lower mortality and a shorter length of hospital stay, making the difference in the survival of hospitalized patients (PATEL A.N. et al., 2020).

The question then became the understanding of how ivermectin acted in the inhibition of the SARS-CoV-2 virus, since as an antiparasitic agent the issue of its antiviral activity was still unknown. In some studies (BOLDESCU et al, 2017; CALY et al, 2020; FRIEMAN et al, 2007; CAO et al., 2020; GREIN et al., 2020; FERNER et al., 2020; CRUMP et al., 2017), ivermectin had been reported as an inhibitor of the nuclear import of viral proteins, as the non-structural protein of the tumor antigen of the ape virus SV40 (an old known molecular biology as cloning vector in ancient techniques of recombinant DNA technology), and also acting in the limitation of infections of other RNA viruses such as viruses of types 1 to 4 of dengue, West Nile, Venezuelan equine encephalitis and influenza. Until, in studies with the HIV-1 virus (human immunodeficiency virus type 1), it was finally associated with the breakdown of the interaction between the ENZYME INTEGRASE of the HIV-1 virus and the heterodimer α / β1 of IMPORTIN, which is the protein responsible for the nuclear import of the INTEGRASE itself.

Since the decade of 1990, the role of integrase as an inhibitor of HIV replication has been suggested by scientists as a promising opportunity in the treatment of viral infections because it is a highly conserved enzyme from an evolutionary point of view and therefore with less genetic variability (SPRINZ, E. 2016). Because it is very conserved, it has greater difficulty in selecting mutations associated with resistance, besides presenting potential synergism with other RNA viruses, including those viruses that had resistance to reverse transcriptase inhibitors. (PURAS L. et al, 1995); (ROBINSON, W.E., 1998) (BEALE K.K., ROBINSON W.E. JR. 2000); (REINKE R, STEFFEN N.R., ROBINSON W.E. JR. 2001)

Although it has been tested in humans for three decades (SMART, T. 1996), its development has been quite “truncated” by the high cost of production and its pharmacokinetic limitations (such as low selectivity due to integrase, difficulties encountered in its injectable use and short half-life time) preventing its clinical use (SPRINZ, E. 2016). However, understanding the evolutionary aspects of this enzyme can help the scientific community understand what possible relationships exist between it and the drugs that interact in its connection with IMPORTIN, especially in the role of destabilization of the import complex that disables viral infections, such as ivermectin. Thinking like this, the team of the Laboratory of Population Genetics and Computational Evolutionary Biology (<u>LaBECom-UNIVISA</u>) designed a study of phylogeny and molecular variance analysis to evaluate the possible levels of genetic diversity and polymorphisms existing in a PopSet of the integrase gene of human immunodeficiency virus 1collected in a Russian population of Kyrgyzstan and available at GENBANK.

## 2. Objective

Evaluate the possible levels of genetic diversity and polymorphisms existing in 153 sequences of the integrase gene of human immunodeficiency virus 1 in the Kyrgyzstan population.

## 3. Methodology

### 3.1. Databank

The 153 gene sequences of the integrase gene of human immunodeficiency virus 1 were collected from GENBANK (https://www.ncbi.nlm.nih.gov/popset/?term=MN888087.1 and participate in a PopSet dipped by Totmenin and collaborators on March 25, 2020 (Popset:1822236350).

### 3.2. Phylogenetics Analyses

For phylogenetic analyses, the previously described nucleotide sequences were used. The sequences were aligned using the MEGA X program (TAMURA et al., 2018) and gaps were extracted for the construction of phylogenetic trees.

### 3.3. Genetic Structuring Analyses

Paired FST estimators were obtained with the software Arlequin v. 3.5 (EXCOFFIER et al., 2005) using 1000 random permutations. The FST matrix generated by the software was used in the construction of a dendrogram based on the UPGMA distance method with the MEGA X software (TAMURA et al., 2018) and the FST and geographic distance matrices were not compared.

## 4. Results

### 4.1. General properties of integrase gene sequences of the HIV-1 human virus

Of the 153 sequences of the gene segment of the integrase gene of human immunodeficiency virus 1 with 882 bp of extension, the analyses revealed the presence of 343 polymorphic sites and of these, 70 sites were parsimoniously informative. The graphical representation of these sites could be seen in a logo built with the WEBLOGO 3 program (CROOKS et al., 2004), where the size of each nucleotide is proportional to its frequency for certain sites. (Figure 1).

**Figure 1:**
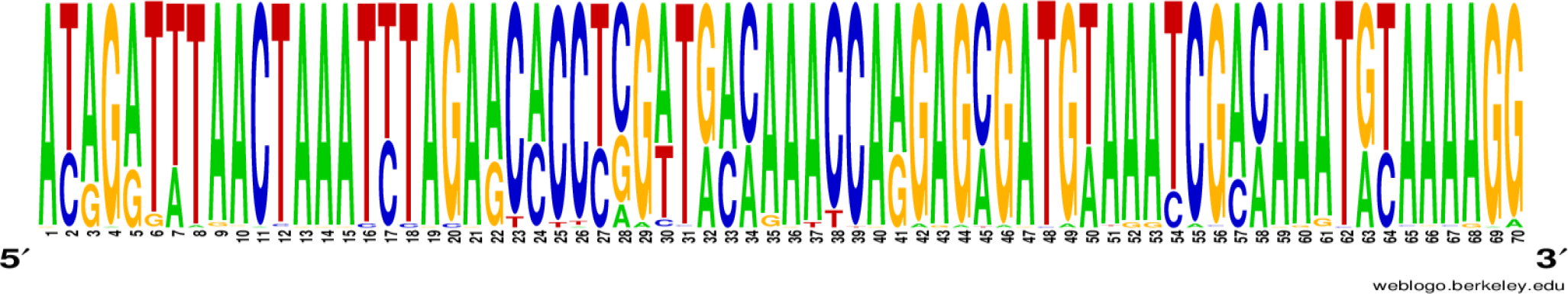
Graphic representation of 70 parsimonious-informative sites of the integrase gene of human immunodeficiency virus 1.

Using the UPGMA method, based on the 70 parsimony-informative sites, it was possible to understand that the 153 haplotypes comprised two distinct groups, here called Bishkek and Osh, in reference to their collection origin and no haplotype sharing was observed between the two groups (Figure 2).

**Figure 2.**
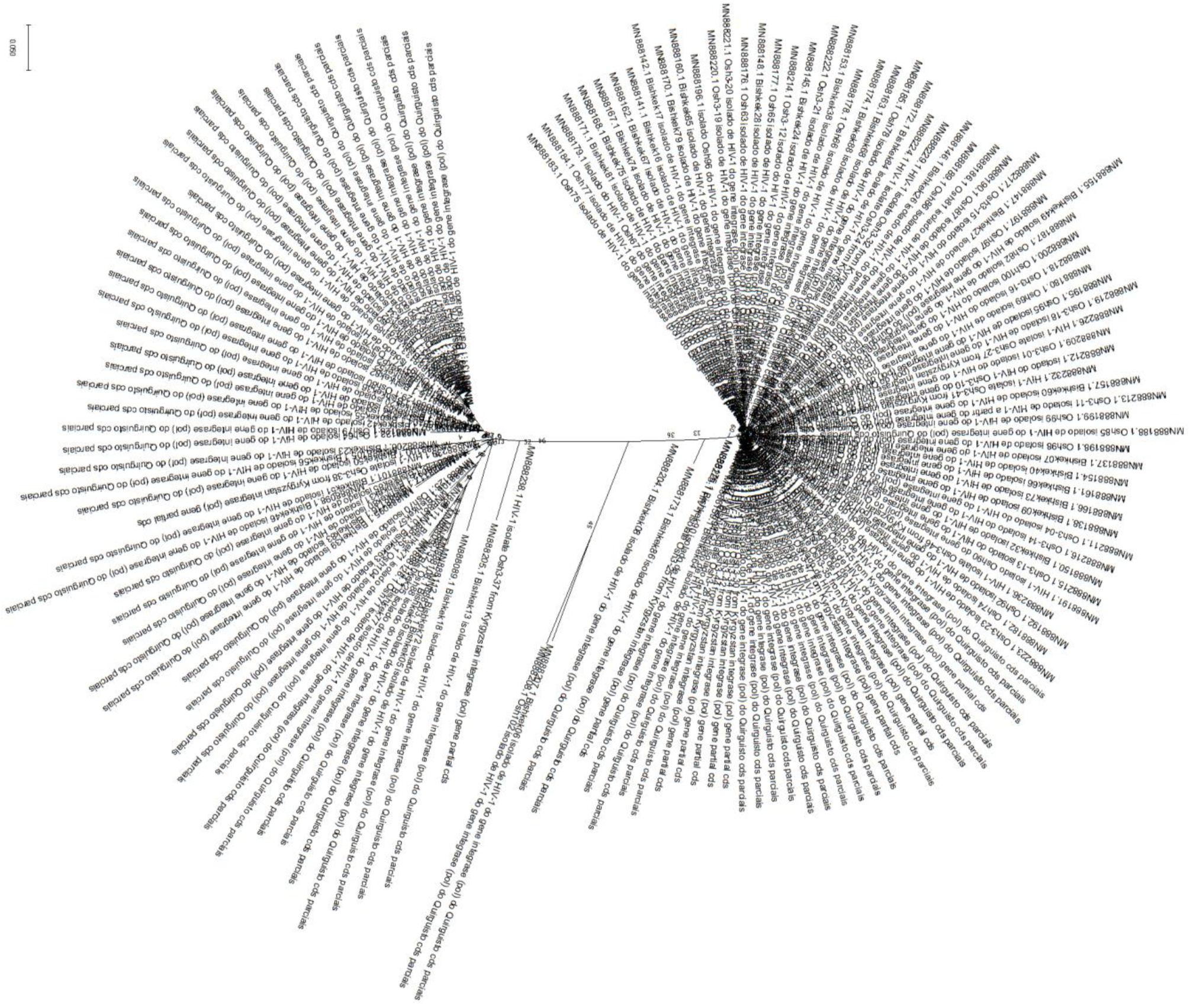
Evolutionary analysis by the maximum likelihood method. The evolutionary history was inferred using the Maximum Likelihood method and the Tamura 3-parameter model [1]. The tree with the highest probability of logging (−1366.35) is shown. The percentage of trees in which the associated taxa group is shown next to the branches. The initial trees for heuristic search were obtained automatically by applying the Join-Join and BioNJ algorithms to an array of distances in estimated pairs using the Tamura 3 parameter model, and then selecting the topology with a higher log probability value. This analysis involved 153 nucleotide sequences. There was a total of 70 positions in the final dataset. Evolutionary analyses were performed on MEGA X.

### 4.2. Genetic Distance Analysis

Analyses based on FST values also confirmed the presence of two distinct genetic entities, with a component of variation greater than 36% and with p value lower than 0.05 with significant evolutionary divergences within the groups (table 1) and also evidenced a high genetic similarity between the sequences that comprised the Oshi group, as well as a greater evolutionary divergence between the sequences that comprised the Bishkek group (table 2); (table 3).

**Table 1.** Paired FST values for the 153 sequences of the integrase gene of human immunodeficiency virus 1 with 882 bp extension.

| Populations | Bishkek | Osh |
| --- | --- | --- |
| Bishkek | 0.00000 | **0.36597** |
| Osh | **0.36597** | 0.00000 |

**Table 2.** Estimates of the mean evolutionary divergence within the groups. Number of base substitutions per location, the average of all sequence pairs within each group.

| Groups | Estimate average | Standard error |
| --- | --- | --- |
| Bishkek | 0,23 | 0,05 |
| Osh | 0,09 | 0,01 |

Standard error estimates are shown above the diagonal. The analyses were performed using the maximum composite likelihood model [1]. This analysis involved 153 nucleotide sequences. All ambiguous positions have been removed for each sequence pair (pair exclusion option). There was a total of 70 positions in the final dataset. Evolutionary analyses were performed on MEGA X.

**Table 3.** Estimates of evolutionary divergence between groups.

| Groups | Bishkek | Osh |
| --- | --- | --- |
| Bishkek | - | **0,27** |
| Osh | **0,27** | - |

The number of base overrides per location of the average of all pairs of sequences between groups is shown. The analyses were performed using the maximum composite likelihood model [1]. This analysis involved 153 nucleotide sequences. All ambiguous positions have been removed for each sequence pair (pair exclusion option). There was a total of 70 positions in the final dataset. Evolutionary analyses were performed on MEGA X.

### 4.3. Molecular Variance Analysis (AMOVA)

Molecular variation analyses of the 153 sequences of the integrase gene of human immunodeficiency virus 1 revealed very significant FST values (FST = 0.36) when analyzed as distinct and even more significant groups when their internal differences were analyzed (in both groups) (Table 4).

**Table 4:** Molecular Variance Analysis, applying Wright’s FST (1969), for the 153 sequences of the integrase gene of human immunodeficiency virus 1 with 882 bp extension

| Variation source | Degrees of freedom. | Sum of squares | Variation components | Percentage of variation. |
| --- | --- | --- | --- | --- |
| **Among the populations** | 1 | 207.72 | 2.65 Va | 36.6% |
| **Within populations** | 151 | 694.62 | 4.60 Vb | 63.4% |
| **TOTAL** | 152 | 902.34 | 7.25 |  |

FST = 0,3659 \*p < 0,05/ Significance tests (1023 permutations). AMOVA design and results: Weir, B.S. and Cockerham, C.C. 1984. Excoffier, L., Smouse, P., and Quattro, J. 1992. Weir, B. S., 1996.

Tau variations (related to the ancestry of the two groups) revealed a significant time of divergence, supported by mismatch analysis of the observed distribution (τ = 44%) and with constant mutation rates between localities (table 5).

**Table 5.** Tau (τ) values for the 153 sequences of the integrase gene of human immunodeficiency virus 1 with 882 bp extension

| **Populations** | **Bishkek** | **Osh** |
| --- | --- | --- |
| Bishkek | 0.00000 | **1.72033** |
| Osh | **1.72033** | 0.00000 |

### 4.4. Molecular diversity analyses

Molecular diversity analyses estimated by θ reflected a significant level of mutations among all haplotypes (transitions and transversions). Indel mutations (insertions or deletions) were not found in either of the two groups studied. The D tests of Tajima and Fs de Fu showed disagreements between the estimates of general θ and π, but with negative and highly significant values, indicating an absence of population expansion. The irregularity index (R= Raggedness) with parametric bootstrap simulated new values θ for before and after a supposed demographic expansion and in this case assumed a value equal to zero for the groups (Table 6); (Table 7).

**Table 6.** Molecular Diversity Indexes for the 153 sequences of the integrase gene of human immunodeficiency virus 1 with 882 bp extension

| **Indexes** | **Bishkek** | **Osh** |
| --- | --- | --- |
| Transitions | 22 | 17 |
| Transversions | 12 | 03 |
| Replacements | 34 | 20 |
| Indels | 0 | 0 |
| $\pi$ | 8.6 | 6.1 |
| $\theta S$ | 8.1 | 5.4 |
| $\theta S$ (d.p) | 2.7 | 2.0 |
| $\theta\pi$ | 8.6 | 6.1 |
| $\theta\pi$ (d.p) | 4.5 | 3.3 |

**Table 7.**
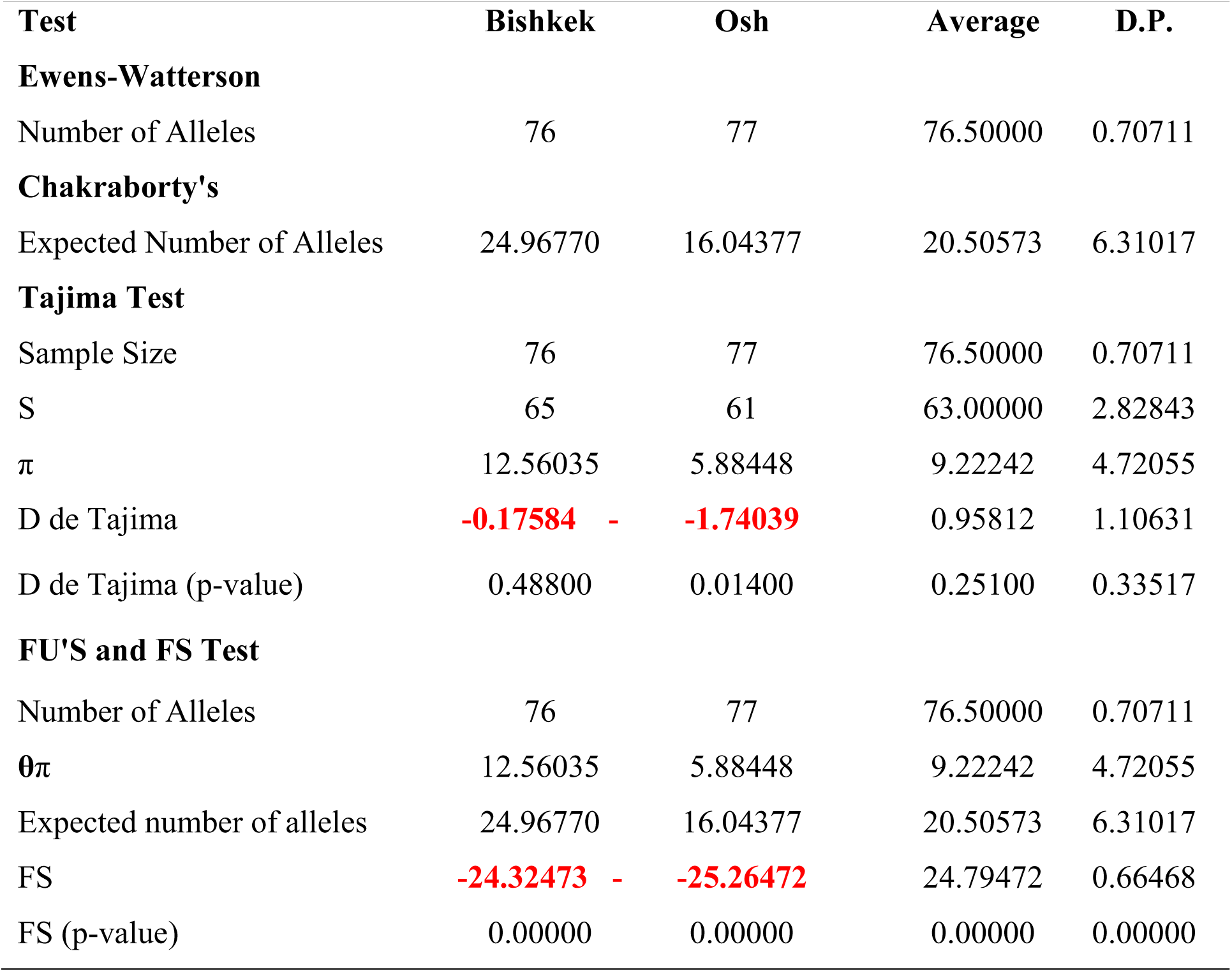
Neutrality tests for the 153 sequences of a segment of the integrase gene of human immunodeficiency virus 1 with 882 bp extension

## 5. Discussion

As the use of phylogenetic analysis and population structure methodologies had not yet been used in this PopSet, in this study it was possible to detect the existence of these two distinct groups for the integrase gene of human immunodeficiency virus 1 in the Kyrgyz region. The groups described here seem to correspond to two HIV-1 subpopulations that co-exist in the same locality and that had their genetic distances supported by FST analyses using the marker in question and its structure sufficiently significant for such interpretation. Different degrees of structuring were detected for each group, being essentially smaller among one of them (Bishkek). These data suggest that the high degree of structuring present in Oshi may be related to a loss of intermediate haplotypes over the generations, possibly associated with an absence of gene flow.

These levels of structuring were also supported by simple phylogenetic pairing methodologies such as UPGMA, which in this case, with a discontinuous pattern of genetic divergence between the groups (supporting the occurrence of geographic undercalculations resulting from past fragmentation events), was observed a large number of branches with many mutational steps. These mutations possibly settled by drift due to the founding effect, which accompanies the dispersal behavior and/or loss of intermediate haplotypes over the generations. The values found for genetic distance support the presence of this discontinuous pattern of divergence between the studied groups, since they considered important the minimum differences between the groups, when the haplotypes between them were exchanged, as well as the inference of values greater than or equal to that observed in the proportion of these permutations, including the p value of the test.

The discrimination of the two genetic entities in the same locality was also perceived when the inter-haplotypic variations were hierarchized in all covariance components: by their intra and interindividual differences or by their intra- and intergroup differences, generating dendrograms that supported the idea that the significant differences found in the Bishkek group, for example, can even be shared in their form, but not in their number, since the result of estimates of the mean evolutionary divergence within the Oshi group were so low.

Since no relationship between genetic distance and geographic distance was made in this study, the lack of gene flow (observed by non-haplotypic sharing) should be supported by the presence of geographic barriers. The estimators θ, although being extremely sensitive to any form of molecular variation (Fu, 1997), supported the uniformity between the results found by all the methodologies employed, and can be interpreted as a phylogenetic confirmation that there is no consensus in the conservation of the gene of human immunodeficiency virus integrase 1 in samples from the same geographical region, being therefore safe to state that the large number of polymorphisms existing, should be reflected, including, in its protein product (integrase enzyme). This consideration provides certainty that an efficient response of drugs that destabilize the integrase-importin link such as ivermectin should not be expected for all HIV1 viruses from humans, whether they come from the same geographic region (as this study shows), or even more from samples from geographically distinct regions and thus, by extrapolating the levels of polymorphism and molecular diversity found in the samples of this study, for other RNA viruses, such as Sars-Cov-2, the integrase-importin relationship may be even more diverse, and may bring less and less functionality to drugs that interact with it in the role of destabilizing the integrase-importin complex, which in turn inhibit or reduce the infectious potential of any RNA virus.

